# Modeling the Interplay between Photosynthesis, CO_2_ Fixation, and the Quinone Pool in a Purple Non-Sulfur Bacterium

**DOI:** 10.1101/704130

**Authors:** Adil Alsiyabi, Cheryl Immethun, Rajib Saha

## Abstract

*Rhodopseudomonas palustris* CGA009 is a purple non-sulfur bacterium (PNSB) that can fix CO_2_ and nitrogen or break down organic compounds for its carbon and nitrogen requirements. Light, inorganic, and organic compounds can all be used for its source of energy. Excess electrons produced during its metabolic processes can be exploited to produce hydrogen gas or biodegradable polyesters (polyhydroxybutyrate). A genome-scale metabolic model of the bacterium was reconstructed to study the interactions between photosynthesis, carbon dioxide fixation, and the redox state of the quinone pool. A comparison of model-predicted flux values with published *in vivo* MFA fluxes resulted in predicted errors of 5-19% across four different growth substrates. The model predicted the presence of an unidentified sink responsible for the oxidation of excess quinols generated by the TCA cycle. Furthermore, light-dependent energy production was found to be highly dependent on the rate of quinol oxidation. Finally, the extent of CO_2_ fixation was predicted to be dependent on the amount of ATP generated through the electron transport cycle, with excess ATP going toward the energy-demanding CBB pathway. Based on this analysis, it is hypothesized that the quinone redox state acts as a feed-forward controller of the CBB pathway, signaling the amount of ATP available.

## Introduction

Purple non-sulfur Bacteria (PNSB) are considered to be among the most metabolically versatile groups of bacteria^1,2^. Within this class, *Rhodopseudomonas palustris* CGA009 (hereafter *R. palustris*) demonstrates this elasticity through its ability to survive in a myriad of diverse environmental conditions^3^. It can grow either aerobically or anaerobically, utilize organic (heterotrophic) or inorganic (autotrophic) carbon sources, and exploit light to obtain energy when growing anaerobically^3^. Several interesting features have been observed in this bacterium, such as its consumption of fatty acids, dicarboxylic acids, and aromatic compounds including lignin monomers^4–6^. It is also one of two known bacteria that can express three unique nitrogenases, each with a different transition-metal cofactor^7^. Furthermore, this metabolically versatile strain’s genome includes the aerobic and anaerobic pathways for three of the four known strategies that microbes use to break down aromatic compounds, such as lignin breakdown products (LBPs)^8^. Harnessing *R. palustris*’ unique metabolic versatilities for the conversion of plant biomass to value-added products, such as polyhydroxybutyrate (PHB)^9^, n-butanol^10^, and hydrogen^11,12^, has garnered increasing interest. However, a system’s level understanding of how the bacterium’s complex web of metabolic modules operates in response to environmental changes is hindering the development of the PNSB as a biochemical chassis.

Several studies conducted on *R. palustris* have shown that in addition to the Calvin-Benson-Bassham (CBB) cycle’s role of carbon assimilation during autotrophic growth, the pathway plays a major role in maintaining redox balance under heterotrophic growth^10,12–14^. It has been shown that heterotrophic growth of the PNSB on substrates that are more reduced than biomass, such as LBPs, is dependent on the availability of an electron sink^13^. CO_2_-fixation using the enzyme ribulose-1,5-biphosphate carboxylase/oxygenase (RuBisCO), nitrogen-fixation through the enzyme nitrogenase^12^, and the supplementation of an electron acceptor (e.g., TMAO)^15^ all prevent the inhibitory accumulation of excess reducing agents. The use of CO_2_ as a redox balancing strategy for the conversion of plant biomass to value-added products is an attractive approach that could increase profitability while improving sustainability. However, the complex interplay between the electrons supplied by the catabolism of different carbon sources, CO_2_ fixation, and the cyclic electron flow during photosynthesis is not fully understood, thus diminishing the ability to engineer this promising bacterium.

A Genome-Scale Metabolic Model (GSMM) provides a mathematical representation of an organism’s metabolic functionalities^16^. The metabolic network represents an organism’s available repertoire of biochemical transformations constructed as a stoichiometric matrix^17^. Due to the underdetermined nature of metabolic networks, optimization tools are used to predict reaction rates for a pre-specified objective function such as the maximization of biomass^18^. One of the most commonly applied optimization tools used to model metabolism is Flux Balance Analysis (FBA). FBA performs a pseudo-steady state mass balance for each metabolite in the network to predict the maximum growth rate and corresponding reaction fluxes during the cell’s exponential growth phase^19^. Due to the high dimensionality of the network, other tools such as Flux Variability Analysis (FVA) are used to determine the sensitivity of growth rate as a function of each reaction flux^20^. Finally, a modified FBA formulation can be used to predict the set of essential genes under a specified growth condition^21^. Thus far, a limited number of small-scale metabolic reconstructions have been developed for PNSB, examining either the central carbon metabolism^22^ or the electron transport chain^23^. However, these models are limited in scope, as they consider less than 4% of the organism’s metabolic functionality and are therefore incapable of capturing system-wide interactions between different metabolic modules. Very recently, a GSMM of the bacterium was reconstructed and used to test an array of cellular objectives during phototrophic growth. Anaerobic growth on acetate, benzoate, and 4-hydroxybenzoate was simulated using 8 different biologically relevant objective functions. The model predicted that the organism primarily optimized for growth, ATP production, and metabolic efficiency. However, the model could further be improved by integrating recently annotated metabolic pathways of lignin monomer degradation^24^, and by making use of experimental data on gene essentiality^25^ and metabolic flux analysis for growth under different carbon sources^13,14^ to validate and refine the network.

In this work, a Genome-Scale Metabolic Model (GSMM) of *R. palustris* was constructed to model the bacterium’s metabolic functionality under different environmental conditions. The model was used to simulate growth under different carbon sources and shows excellent agreement with experimentally measured fluxes^13,14^. Gene essentiality analysis was also performed for aerobic and anaerobic growth on acetate. The predicted essential genes were compared with available trans-mutagenesis data^25^, and an accuracy of 84% was achieved. After the model indicated the presence of an unidentified quinol sink, *in silico* simulations were combined with published *in vivo* flux measurements^13,14^ to study the effect (and extent) of the quinone redox state on cellular growth, electron transport rate, and CO_2_ fixation. These results suggest that redox state acts as a feed-forward controller of the highly energy-demanding CBB cycle by regulating the rate of light-generated ATP. Overall, an understanding of the metabolic control points of this interconnected system constitutes the first step towards engineering strains capable of more efficiently harnessing photosynthetic energy and rerouting this energy towards bio-production and lignin valorization.

## Methods

### Model reconstruction

A draft model was first generated in KBase^26^ based on *R. palustris*’ genome (downloaded from the NCBI database on 04/12/2018). KBase uses annotated features in the genome to construct a list of reactions associated with genes in the organism. Previously published work of the bacterium’s metabolic network^22^ was used to manually curate pathways from the central carbon metabolism and to ensure correct cofactor usage and gene association. This resulted in an expanded network of high-confidence reactions, all associated with genes in *R. palustris*. Experimentally measured concentrations of biomass components are available for *R. palustris* when grown on acetate^13^ and were used to develop the biomass equation (see supplementary file 1). To minimize the addition of low-confidence reactions during gapfilling, the process was broken down into two steps. First, a subset of high-confidence reactions from a recently published genome-scale model of *R. palustris^27^* was added to the draft model. Here, high-confidence reactions are those that are associated with a published source of annotation. At this point, the majority of the reactions required to gapfill the biomass equation existed in partially incomplete linear pathways. Therefore, the ModelSEED database^28^ was used to gapfill the model generated in KBase. In addition, annotated metabolic pathways for the breakdown of multiple aromatic compounds including lignin breakdown products were found in literature^24^ and in organism-specific biochemical databases^29,30^, and were subsequently added to the model (supplementary file 2). Finally, annotated *R. palustris* genes were mined from three databases (KEGG^29^, BioCyc^30^, and UniProt^31^) to validate the Gene-Protein-Reaction (GPR) associations established in the model and to include GPR relationships for reactions added during the gapfilling process (supplementary file 3).

### Model simulations

Parsimonious Flux Balance Analysis (pFBA)^32^ was used to simulate growth under different environmental conditions. pFBA is analogous to FBA but adds an outer objective that minimizes the sum of all reaction fluxes. Objective tilting^33^ was used to formulate both objectives in one function as shown below.

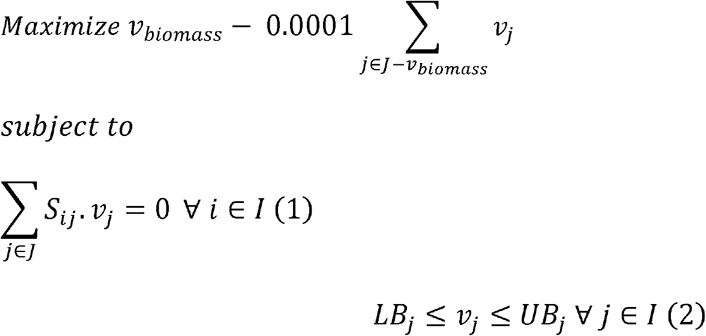

Where *I* and *J* are the sets of metabolites and reactions in the model, respectively. *S*_*ij*_ is the stoichiometric coefficient of metabolite *i* in reaction *j* and *v*_*j*_ is the flux value of reaction *j*. Parameters *LB*_*j*_ and *UB*_*j*_ denote the minimum and maximum allowable fluxes for reaction *j*, respectively. *v*_*biomass*_ is the flux of the biomass reaction which mimics the cellular growth rate.

### Model validation

Metabolic Flux Analysis (MFA) measurements from anaerobic growth on acetate^13^, fumarate, succinate, and butyrate^14^ were compared with model predicted fluxes. Model accuracy for each growth condition was calculated by taking the sum of percent errors between pFBA predicted and MFA values (see supplementary file 4 for example). In addition, *R. palustris’* essential genes, determined experimentally for aerobic growth on acetate^25^, was used to validate the essential genes predicted by the model. Gene essentiality was predicted in the model by sequentially knocking out each reaction and determining the resulting effect on the biomass reaction rate^21^. If a reaction knockout resulted in a predicted growth rate that was less than 10% of the wild type growth rate, the reaction was considered essential. Reaction GPRs were then used to map the list of essential reactions to essential genes. Finally, the list of experimentally determined essential metabolic genes^25^ were compared with model predicted essential genes to determine the specificity and sensitivity of the predictions (see supplementary file 5).

## Results and Discussion

### Model Reconstruction and validation

A summary of the model’s major statistics is shown in Figure 1A. Overall, the 940 genes associated with model reactions account for 62% of the genes involved in energy metabolism, biosynthesis, carbon & nitrogen metabolism, and cellular processes in *R. palustris*’ genome^3^. Experimental measurements of biomass component concentrations were obtained for growth on acetate^13^ (Figure 1B) and converted into stoichiometric coefficients for the model’s biomass equation (see supplementary file 1). Thus, an initial high-confidence model containing 540 genes and 915 reactions with no orphan reactions was constructed. The gap-filling procedure was carried out next in KBase^26^ using reactions from the ModelSEED database^28^. Organism-specific biochemical databases including KEGG, UniProt, and BioCyc were then used to annotate added reactions (Figure 1C). This resulted in the addition of 328 annotated and 110 unannotated (orphan) reactions. The inclusion of these reactions is necessary to ensure biomass production. pFBA was used to simulate growth on a number of different carbon sources, including carboxylic acids (acetate, fumarate, succinate and butyrate) and lignin monomers (supplementary file 2). pFBA is analogous to FBA but adds an outer objective that minimizes the sum of all reaction fluxes (see Methods). This is justified by the assumption that cells synthesize the minimum amount of cellular machinery required to maintain the maximal growth rate^32^. Simulating growth using pFBA has two main advantages over FBA. First, pFBA avoids unrealistic flux predictions for reactions participating in thermodynamically infeasible cycles (TICs)^34^. TICs are usually removed from GSMs to avoid false predictions; however, when analyzing highly connected networks like that of *R. palustris*, removing these cycles can lead to the model missing certain functionalities and metabolic modes utilized by the organism. pFBA avoids these false predictions by the additional constraint that reaction fluxes should be minimized. Second, the pFBA formulation results in a significantly reduced set of optimal solutions compared to FBA. Flux Balance Analysis usually results in a large number of alternate optimal solutions (especially in highly connected networks), most of which are not biologically relevant, and can therefore lead to false conclusions^35^. pFBA’s additional objective greatly restricts the solution space and leads to more biologically insightful conclusions^32^. Essentiality analysis identified 368 essential reactions, out of which 249 were associated with gene annotations in the model. Comparison with *in vivo* gene essentiality data for aerobic growth on acetate^25^ was then used to check the model accuracy (Figure 1D). The calculated sensitivity and false negative rate (FNR) are consistent with recently published GSMMs^36,37^. Moreover, given that this is a non-model organism with no well-characterized close relatives, high-confidence annotation was not available for less-studied pathways, therefore an automated pipeline like GrowMatch^38^ could not be implemented to further improve essentiality predictions.

**Figure 1.**
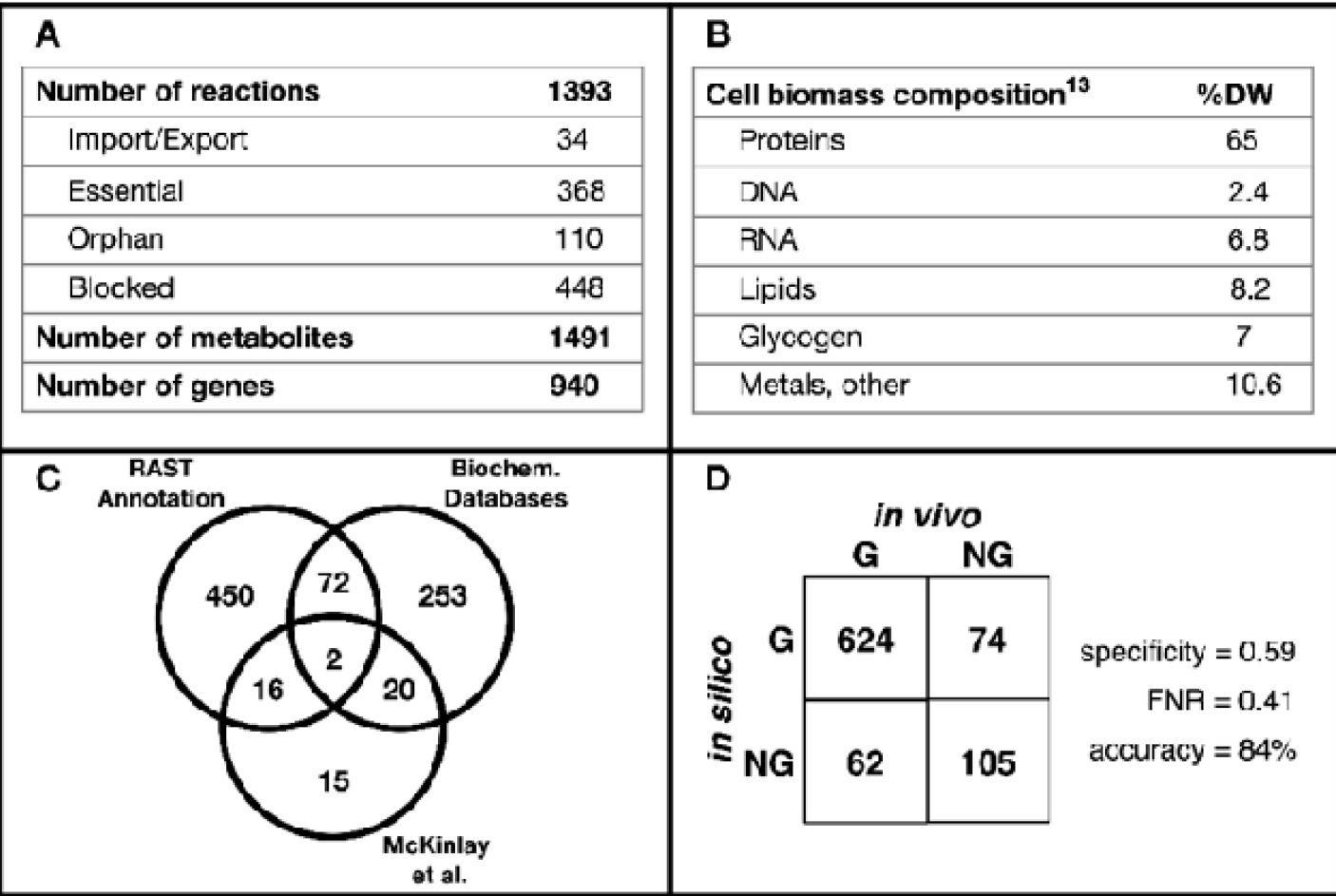
Summary of model statistics and validation. **(A)** Overall model statistics. **(B)** Model biomass component compositions. **(C)** Sources of gene annotation. **(C)** Gene essentiality analysis results.

### The effect of the quinone pool on light uptake, carbon dioxide fixation, and growth

During initial growth simulations, growth was observed to be hindered due to the accumulation of excess quinols formed in the TCA cycle. Since no high-confidence reaction was found to consume quinols in *R. palustris*, a quinol “sink” reaction was added to the model. Sink reactions are often incorporated into metabolic models when a metabolite is known to be produced during metabolism but for which no means of consumption have been identified^39^, or to describe the accumulation of a storage compound^39^ (e.g. glycogen). Furthermore, recent experimental work with *R. palustris* TIE-1 reported the presence of an unidentified quinol-oxidizing reaction that had not been accounted for previously^40^, giving further support to this prediction. To determine the effect of the quinone pool on growth, pFBA simulations were conducted under different quinol sink rates to qualitatively predict how changes in the quinone redox state affected the rest of the metabolic network. The quinol sink reaction was treated as a parameter in the model and pFBA simulations were conducted at varying quinol oxidation (sink) rates to determine how light uptake (i.e. Electron Transport Rate or ETR), growth, and CO_2_ fixation are affected by changes in the quinone redox state (Figure 2). Carbon uptake was restricted to a maximum value of 100 mmol/gDW/hr for acetate and 50 mmol/gDW/hr for fumarate, succinate, and butyrate to ensure the same number of carbons were being up taken. MFA values were scaled to the same carbon uptake rates^13,14^. For growth on butyrate, the supplementation of CO_2_ is required for growth, as the substrate is more reduced than biomass and requires an electron sink^14^. The media was supplied with CO_2_ at a maximum uptake rate of 32.1 mmol/gDW/hr to match MFA observations. Since steady-state GSMMs cannot capture metabolite concentrations, the redox state cannot be calculated directly. Instead, the qualitative behavior of the redox state was predicted by varying the rate of the quinol sink. As the quinol oxidation rate increases, the quinone pool becomes more oxidized. Using experimental MFA data^13,14^, the quinol oxidation rate was predicted for each of the four substrates (Table 1). These values were calculated by minimizing the sum of errors between the *in silico* generated pFBA fluxes and the *in vivo* MFA flux values. The table also shows the quinone reduction rate through the TCA cycle for each carbon source. The percentage of CO_2_ fixed was defined as the rate of CO_2_ fixation divided by the total rate of CO_2_ produced metabolically. Figure 3 shows the resulting flux predictions obtained at the predicted quinol oxidation rates for growth on acetate (figure 3A), and the calculated percent errors of these predictions for each carbon substrate (figure 3B). A comparison of flux predictions with MFA values for the other three carbon sources is provided in supplementary file 2.

**Figure 2.**
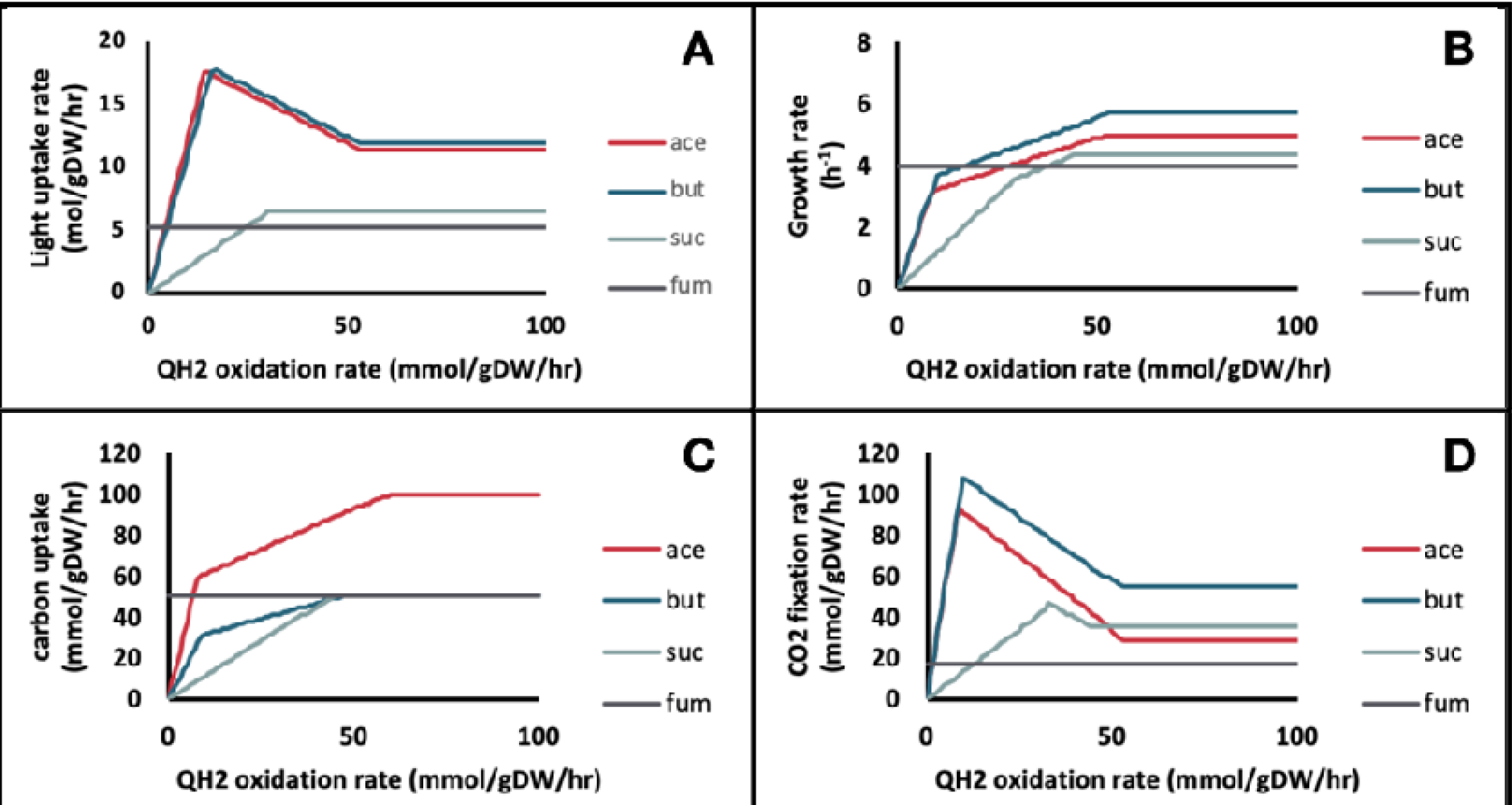
Effect of the Quinol sink rate on: (A) Light uptake rate, (B) Growth rate, (C) Carbon source.

**Table 1.** Predicted reaction rates for growth on four different carbon sources

| Carbon source | QH<sub>2</sub> oxidation rate (mmol/gDW/hr) | Q reduction rate<sup>a</sup> (mmol/gDW/hr) | Electron transport rate (mol/gDW/hr) | CO<sub>2</sub> fixation rate (mol/gDW/hr) | % CO<sub>2</sub> fixed<sup>b</sup> | Net CO<sub>2</sub> excretion rate (mmol/gDW/hr) |
| --- | --- | --- | --- | --- | --- | --- |
| Acetate | 52.5 | 39.1 | 5.3 | 29.7 | 73.2 | 10.9 |
| Butyrate | 54.9 | 37.4 | 5.4 | 57.8 | - | -18.6<sup>c</sup> |
| Succinate | 49.8 | 49.2 | 3.0 | 35.6 | 50.6 | 34.8 |
| Fumarate | 0 | 0 | 2.3 | 17.3 | 25.1 | 51.5 |

<sup>a</sup>The rate of quinone reduction in the TCA cycle.
<sup>b</sup>The rate of CO<sub>2</sub> fixation divided by the rate of total CO<sub>2</sub> produced.
<sup>c</sup>CO<sub>2</sub> was supplied in the media during growth on butyrate.

**Figure 3.**
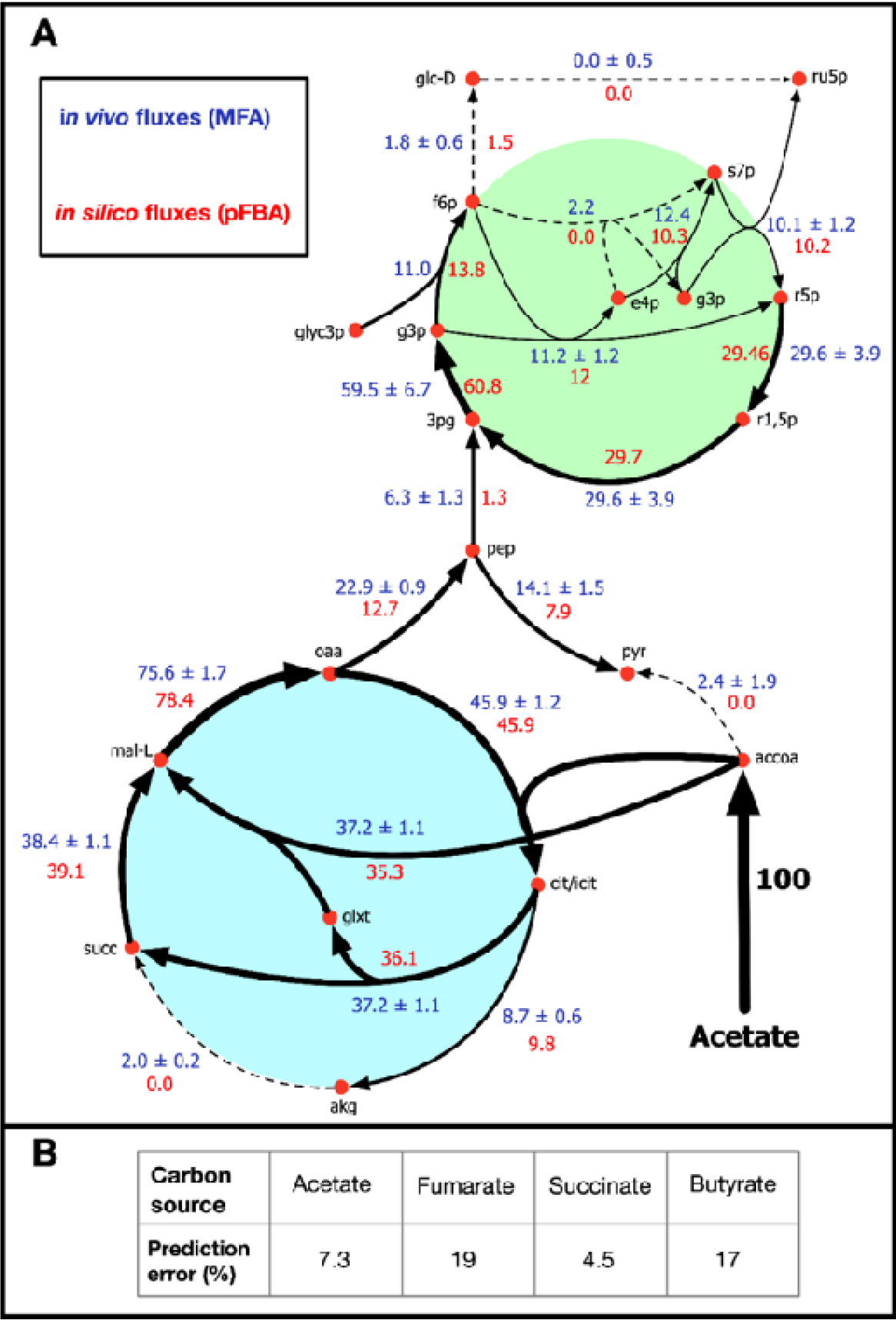
Comparison of model-predicted vs MFA-generated flux values for reactions involved in central carbon metabolism. **(A)** Metabolic flux map showing reaction rates for growth on acetate **(B)** Percentage error

For growth on acetate and butyrate, light uptake (i.e. ETR) shows two distinct regions based on the extent of quinol oxidation (Figure 2A). Under low oxidation rates, flux through the quinol-producing succinate dehydrogenase reaction is avoided by using the glyoxylate shunt and subsequently the CBB cycle. Therefore, both light uptake and CO_2_ fixation increase rapidly in this region. In the second region, at high quinol oxidation rates, flux shifts toward the oxidative TCA cycle. Therefore, in this region, both the Electron Transport Chain (ETC) activity and the rate of CO_2_ fixation decrease with increasing quinol oxidation. Furthermore, as can be seen from Table 1, the ratio of quinol oxidation rate to quinone reduction rate is similar for both carbon sources. Due to the supplementation of CO_2_ during growth on butyrate, the percentage of CO_2_ fixation could not be calculated. During growth on succinate, the production of quinols through succinate dehydrogenase cannot be avoided, therefore light uptake rate increases linearly with the rate of quinol oxidation. Moreover, the rates of quinol oxidation and quinone reduction are equivalent, indicating that the quinone pool is more reduced when compared to the redox state during growth on acetate and butyrate. This leads to a reduced electron flow through the ETC, and subsequently lower ATP generation. Finally, the model predicts that during growth on the highly oxidized (compared to cell biomass) carbon source fumarate, the rate of the quinol sink does not affect the flux distribution.

A similar parameter sampling procedure was performed to determine the effect of light uptake on growth. Light uptake rate was set as a parameter and the quinol oxidation rate was fixed to the value predicted based on MFA fluxes (Figure 4). Again, the plots show two distinct growth regions: (i) a low-light (LL) energy-limited region, and (ii) a high-light (HL) carbon-limited region. In the LL region, growth is highly dependent on the amount of light available and the model predicts that all of the ATP produced is used to convert the carbon source into biomass precursors. Therefore, no ATP remains for the energy-intensive CBB pathway. In the HL region, the maximum substrate uptake rate is reached, and the carbon source cannot be incorporated any faster. The additional energy produced from light is then directed towards CO_2_ fixation. Although the model predicts that the rate of CO_2_ fixation increases linearly with light uptake rate, kinetic and thermodynamic constrains on the highly inefficient CO_2_-fixing RuBisCO enzyme^41^ hinder this process at high light uptake.

**Figure 4.**
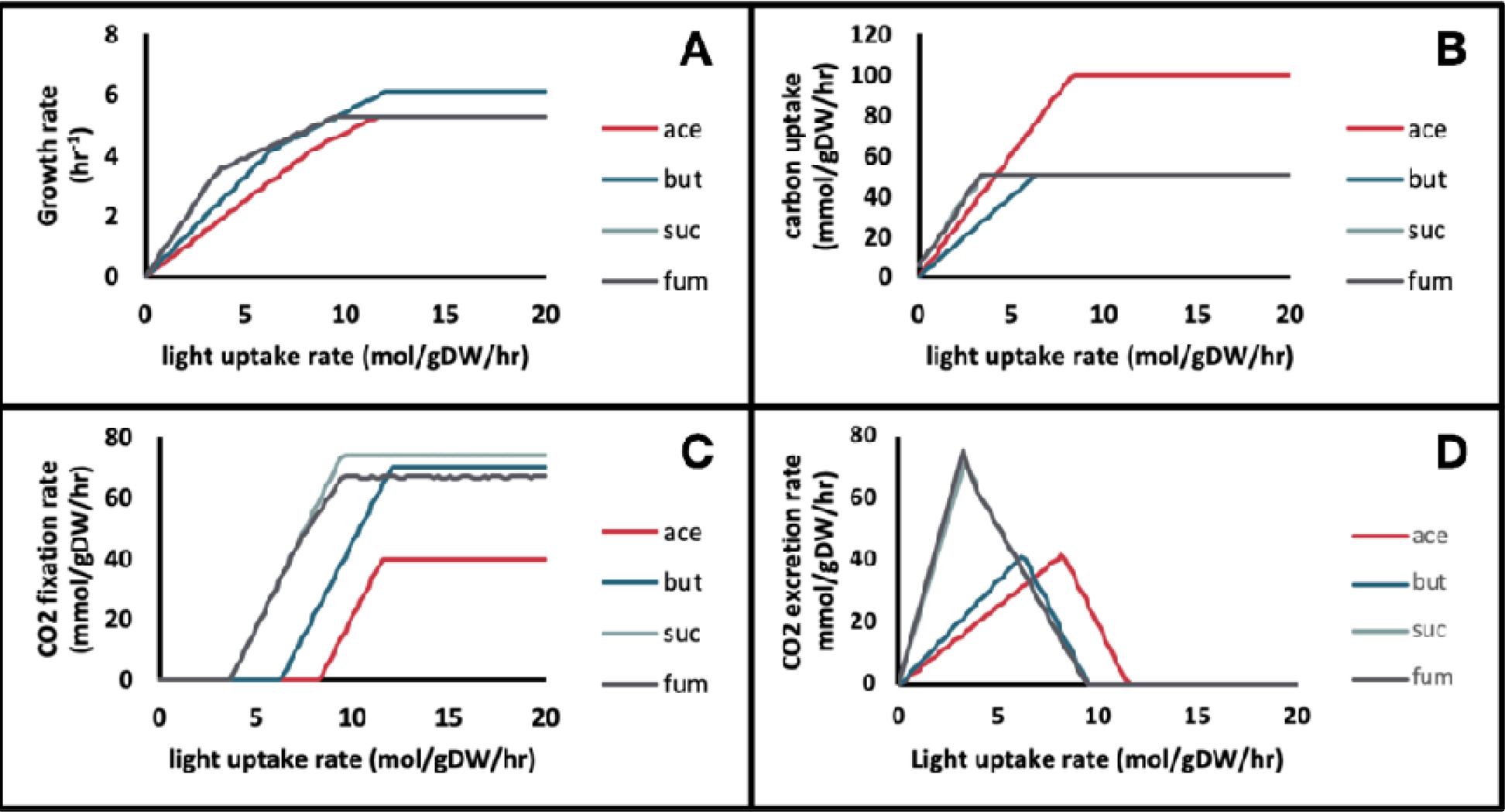
Effect of the light sink rate on (A) Growth rate, (B) Carbon source uptake rate, (C) Carbon fixation rate, and (D) Carbon dioxide excretion rate for growth on four carbon sources. Ace: acetate, but: butyrate, suc: succinate, fum: fumarate. In A, B, and D, the lines for succinate and fumarate lie on top of each other.

### Proposed mechanism for the interplay between the quinone redox state, the electron transport rate, and CO_2_ fixation

Based on the effect of the quinol oxidation rate on light uptake and on the model’s flux distribution, a mechanistic explanation of the system-wide metabolic interactions can be postulated. As shown in Figure 5, increased flux through the oxidative TCA cycle leads to the accumulation of reduced quinols. This in turn leads to a restriction in the flow of electrons through the ETC and consequently in the amount of ATP produced. The CBB system thus lacks the energy required to fix CO_2_. Therefore, the quinone redox state is predicted to act as a feed-forward controller to the energetically expensive CBB pathway, indicating how much ATP is available at a given condition.

**Figure 5.**
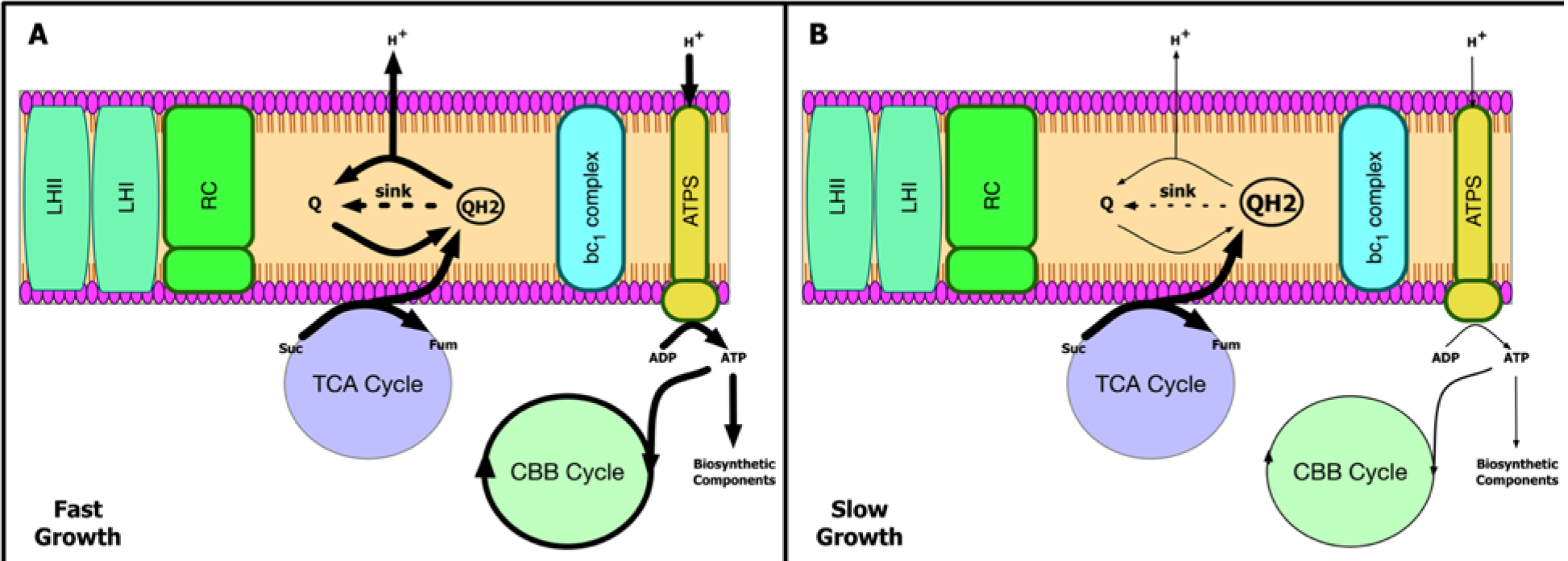
Schematic of a proposed mechanism for the interaction between the quinone redox state, electron transport rate, and carbon fixation. **(A)** High rate of quinol oxidation. **(B)** Low rate of quinol

Comparison of pFBA generated growth simulations with MFA data leads to the hypothesis that an unidentified quinone:oxidoreductase reaction has to occur to obtain the observed flux distribution. A previous study on the PNSB *R. capsulatus* suggests that complex I, the NADH:quinone oxidoreductase enzyme, is responsible for the observed quinol oxidation through reverse electron flow^42^. However, the model predicts that the rate of quinol oxidation required cannot be accounted for through complex I only, which showed low activity. Furthermore, based on the high thermodynamic cost of reverse electron flow, it appears unlikely that it can account for the predicted rate of quinol oxidation^23^.

Although the source of quinol oxidation (sink) has yet to be identified, there are a number of candidate reactions that can carry out this role. Primarily, the malate:quinone dehydrogenase (MDH) appears to be a potential reaction for oxidizing excess quinols. In the forward direction, this reaction converts malate into oxaloacetate and produces ubiquinol in the process. A second NAD-dependent malate dehydrogenase is also coded for by *R. palustris* that could perform the same function. Knocking out and over-expressing these enzymes could be employed to investigate their role in ETR, ATP production, and CO_2_ fixation.

## Conclusion

In this study, a genome-scale metabolic network was used to propose a system-wide mechanistic model of the interactive system that includes photosynthesis, carbon dioxide fixation, and the quinone redox state. The model was validated using experimental genome essentiality data^25^ (84% accuracy), and flux measurement data^13,14^ on four carbon sources (5-19% prediction error). Model simulations predict the presence of an unidentified quinol sink. Predictions also indicate that the extent of CO_2_ fixation is dependent on the amount of ATP present, with the quinone redox state acting as a feed forward signal to the CBB system. Going forward, the proposed mechanism can be used to generate strategies for engineering strains capable of more efficiently harnessing photosynthetic energy, and that have the ability to reroute energy towards bioproduction and lignin valorization. Future experimental work will be conducted to measure the electron transport rate, intracellular ATP concentration, and RuBisCO gene expression across different quinone redox states to strengthen the proposed hypothesis and further refine the model.

## Acknowledgement

Funding to support this work was provided by University of Nebraska-Lincoln Faculty Startup Grant to Rajib Saha. We thank all members of the Systems and Synthetic Biology Laboratory for collegial discussions.

## Author contributions

A.A., C.I., and R.S. conceived the idea of the project. A.A. reconstructed the model and conducted computational analysis. All authors designed research, wrote and edited the manuscript.

## Data Availability

All data generated or analyzed during this study are included in this published article (and its Supplementary Information files).

## Supplementary information

1. Supplementary file 1: Biomass equation components
2. Supplementary file 2: Lignin monomer breakdown pathways and flux predictions for growth on butyrate, succinate, and fumarate.
3. Supplementary file 3: Model files for *R. palustris*
4. Supplementary file 4: Model flux predictions accuracy calculations
5. Supplementary file 5: Gene essentiality analysis results

